# ERECTA family signaling controls cell fate specification during ovule initiation in Arabidopsis

**DOI:** 10.1101/2024.10.10.617662

**Authors:** Alex M. Overholt, Christina Elaine Pierce, Calen Seth Paleologos, Elena D. Shpak

## Abstract

The number of seeds produced by a plant depends on the number of ovules initiated within the carpels by a meristematic tissue, the placenta. Previous studies indicated that ERECTA-LIKE1 (ERL1) and ERL2 receptors and their extracellular ligand EPIDERMAL PATTERNING FACTOR-LIKE2 (EPFL2) promote seed density while ERECTA inhibits it. Here, we show that all three receptors and two ligands, EPFL1 and EPFL2, promote ovule initiation. After initiation, EPFL1 and EPFL2 redundantly regulate the elongation of nascent ovules and promote the growth of the funiculus. The synergy of EPFL1 and EPFL2 with *CUP-SHAPED COTYLEDON* boundary genes in the regulation of ovule number and shape suggests cooperation in their function. Ovule initiation starts with the specification of ovule founder cells that are separated by boundary cells. During ovule bulging, a new cell type, characterized by a high proliferative rate, forms between ovule founder and boundary cells. In the *epfl1 epfl2* mutant, marker genes for ovule founder and boundary cells are expressed much more broadly. Our data indicate that ERECTA family signaling is necessary for efficient cell fate separation at the earliest stages of ovule initiation. Later, it promotes the formation of a zone of high proliferation at the base of the ovule.

## Introduction

Plants carefully regulate lateral organ initiation to achieve specific organ spacing and overall morphology. The number of seeds produced by a flower depends on the number of ovules initiated by the placenta, a meristematic tissue located inside carpels. From an evolutionary standpoint, carpels are modified leaves with fused margins. Each margin gives rise to the carpel margin meristem, a section of which differentiates into the placenta. In *Arabidopsis thaliana*, a pistil is made of two fused carpels and contains four strands of the placenta. Ovules initiate from the placenta starting from stage 9 of floral development, after the fusion of carpels but before the complete closure of the pistil on the top (Smyth et al., 1990; Bowman et al., 1991). This process is asynchronous as the placenta continues to elongate, and new ovules can form between existing ones (Yu et al., 2020). Overall, the number of ovules depends on the length of the placenta and ovule spacing. Ovule development has been extensively characterized and divided into stages 1-I through 3-VI (Robinson-Beers et al., 1992; Schneitz et al., 1995; Sieber et al., 2004; Vijayan et al., 2021). During initiation, ovules resemble finger-like protrusions (stages 1-I through 2-II). During stage 2-III, the base of the ovule becomes the stalk-like funiculus, the middle domain called the chalaza initiates the bowl-shaped integuments, and, on the top, the megaspore mother cell develops within the nucellus.

The initiation of ovules, in many ways, resembles the initiation of above-ground lateral organs, all initiated by meristematic tissues. Accumulation of the plant hormone auxin in the epidermis of cotyledons, leaves, and floral organs is one of the first signs of lateral organ initiation (Benková et al., 2003; Friml et al., 2003). Polar transport of auxin into maxima is essential for radial positioning of these organs (Reinhardt et al., 2000; Reinhardt et al., 2003; Lampugnani et al., 2013). Similarly, ovule initiation depends on the formation of auxin maxima; plants that cannot transport auxin or sense it produce significantly fewer ovules (Benková et al., 2003; Bencivenga et al., 2012; Cucinotta et al., 2021). In addition to auxin, several other phytohormones regulate the number of ovules in a carpel. The loss of enzymes that degrade cytokinins leads to an increase in ovule number, while the inability to sense cytokinins due to mutations in their receptors decreases ovule number (Bartrina et al., 2011; Bencivenga et al., 2012). It has been proposed that cytokinins regulate ovule initiation by increasing the length of the placenta and by promoting polar auxin transport (Bencivenga et al., 2012; Galbiati et al., 2013; Cucinotta et al., 2016). Reduced brassinosteroid signaling decreases ovule number, while increased brassinosteroid responses enlarge it (Huang et al., 2013). Similar to cytokinins, brassinosteroids regulate placenta length and polar auxin transport (Huang et al., 2013; Hu et al., 2022). Furthermore, these two hormones function synergistically in the regulation of ovule initiation (Zu et al., 2022). While cytokinins and brassinosteroids increase ovule number, gibberellins decrease it and they do that independently of polar auxin transport (Gomez et al., 2018).

Several transcription factors regulate ovule initiation and spacing. The AP2 domain transcription factor AINTEGUMENTA (ANT) is expressed early in organ primordia and has been shown to promote the initiation of ovules and floral organs (Elliott et al., 1996; Klucher et al., 1996). CUP SHAPED COTYLEDON (CUC) transcription factors, CUC1, CUC2, and CUC3, are expressed in the boundaries of aboveground organ primordia (Ishida et al., 2000; Takada et al., 2001; Vroemen et al., 2003; Gonçalves et al., 2015). They regulate the separation of cotyledons, flowers, floral organs, and the spacing of ovules (Aida et al., 1997; Ishida et al., 2000; Vroemen et al., 2003; Galbiati et al., 2013; Gonçalves et al., 2015). Genetic and molecular analyses suggest that gibberellins function upstream of CUC2 in the regulation of ovule initiation (Barro-Trastoy et al., 2020).

In this study, we explore the role of ERECTA family (ERf) receptor kinases and their ligands in ovule initiation. The ERf receptors (ERfs) are localized in the plasma membrane where they sense small extracellular cysteine-rich proteins from the *EPIDERMAL PATTERNING FACTOR (EPF)/EPF-like (EPFL)* gene family (Hara et al., 2007; Lee et al., 2012). In Arabidopsis, there are three *ERf* genes: *ERECTA (ER), ERECTA-LIKE1 (ERL1),* and *ERL2* (Shpak et al., 2004), and eleven *EPF/EPFL* genes (Hara et al., 2009). Most EPF/EPFL ligands have been shown to function as agonists, but STOMAGEN/EPFL9 is an antagonist (Sugano et al., 2010; Ohki et al., 2011; Lee et al., 2015). Which ligands are able to activate the receptors depends on the presence of a co-receptor TOO MANY MOUTHS (TMM) that is constitutively bound to ERfs in selected tissues (Lin et al., 2017). SOMATIC EMBRYOGENESIS RECEPTOR-LIKE KINASES (SERKs) heterodimerize with ERfs, presumably only in the presence of EPF/EPFL ligands (Meng et al., 2015). This signaling pathway regulates multiple aspects of development, including stomata formation, elongation of aboveground organs, shoot apical meristem size, leaf initiation, and floral organ structure (Torii et al., 1996; Shpak et al., 2004; Shpak et al., 2005; Pillitteri et al., 2007a; Abrash et al., 2011; Bemis et al., 2013; Chen et al., 2013; Shpak, 2013; Uchida et al., 2013; Chen and Shpak, 2014; Kosentka et al., 2019; Jiang et al., 2022). Previously, a study of seed density using natural accessions of Arabidopsis examined the role of ERfs and their ligand EPFL2 in seed density and ovule spacing (Kawamoto et al., 2020). That study showed that the *epfl2* mutant produces significantly fewer seeds compared to the wild type, and ovules are irregularly spaced. A reporter gene assay indicated that *EPFL2* is expressed in the boundaries between ovules. Analysis of mutants indicated that *ERL1* and *ERL2* increase seed density while *ER* decreases it. Seed density depends on ovule spacing during initiation and the following elongation of carpels. For example, an increased seed density in the *er* mutant can be caused either by increased ovule initiation or by decreased ability of a carpel to elongate. The previous study did not investigate these two processes individually.

To gain insight into the role of ERf and EPFLs in ovule initiation, we analyzed ovule density in a range of mutants during floral stage 10, shortly after ovules are initiated. Our analysis indicates that two ligands, EPFL1 and EPFL2, synergistically promote initiation and elongation of ovule primordia during that stage. It also uncovered a positive role of all three ERfs in ovule initiation. The genetic interactions also revealed that *EPFL1* and *EPFL2* regulate ovule spacing synergistically with *CUC3* and ovule elongation synergistically with *CUC2* and *CUC3*. We analyzed the expression of transcription factors *DORNROSCHEN/ENHANCER OF SHOOT REGENERATION 1 (DRN/EST1)* as the earliest marker of ovule initiation (Kirch et al., 2003; Kawamoto et al., 2020) and *CUC3* as the earliest marker of spaces between ovules (Yu et al., 2020). These experiments uncovered that EPFL1 and EPFL2 are necessary to reinforce cell fate identity in the placenta. Once an ovule is initiated, EPFL1 and EPFL2 promote cell proliferation at its base.

## Results

### EPFL1 and EPFL2 regulate ovule density in a partially redundant manner

Previous research demonstrated a role for *EPFL2* in the positive regulation of seed density and control of ovule spacing (Kawamoto et al., 2020). To investigate how *EPFL* genes contribute to ovule initiation, we screened *epfl* single and higher-order mutants. Measurements of ovule number and placenta length for stage 10 flowers, when most ovules are formed (Yu et al., 2020), were taken and ovule density was calculated by dividing ovule number by placenta length. The *epfl2* single mutant showed significantly reduced ovule density (Fig. 1A and B). While the ovule density of *epfl1* showed no change, the *epfl1 epfl2* double mutant (named *epfl1,2* in figures) had a significant decrease in ovule number and density relative to *epfl2* without altering placenta length (Fig. 1A and B, Fig. S1). The reduction in ovule density in *epfl1 epfl2* leads to broader boundary regions between ovule primordia (Fig. 1A). These data suggest that *EPFL1* and *EPFL2* synergistically promote ovule initiation. The simultaneous loss of *EPFL4* and *EPFL6* did not affect ovule density or placental length (Fig. S2). The quadruple *epfl1 epfl2 epfl4 epfl6* mutant had a significant decrease in the placenta length but no change in the ovule density relative to *epfl1 epfl2* (Fig. S2). These data indicate that EPFL4 and EPFL6 do not promote ovule initiation, and all four ligands synergistically promote placenta elongation during early flower development. We did not detect a significant decrease in the ovule density in *er erl1 erl2* versus *epfl1 epfl2* (Fig. S2), suggesting that, out of all EPFLs, EPFL1 and EPFL2 are the only positive regulators of ovule initiation.

**Figure 1.**
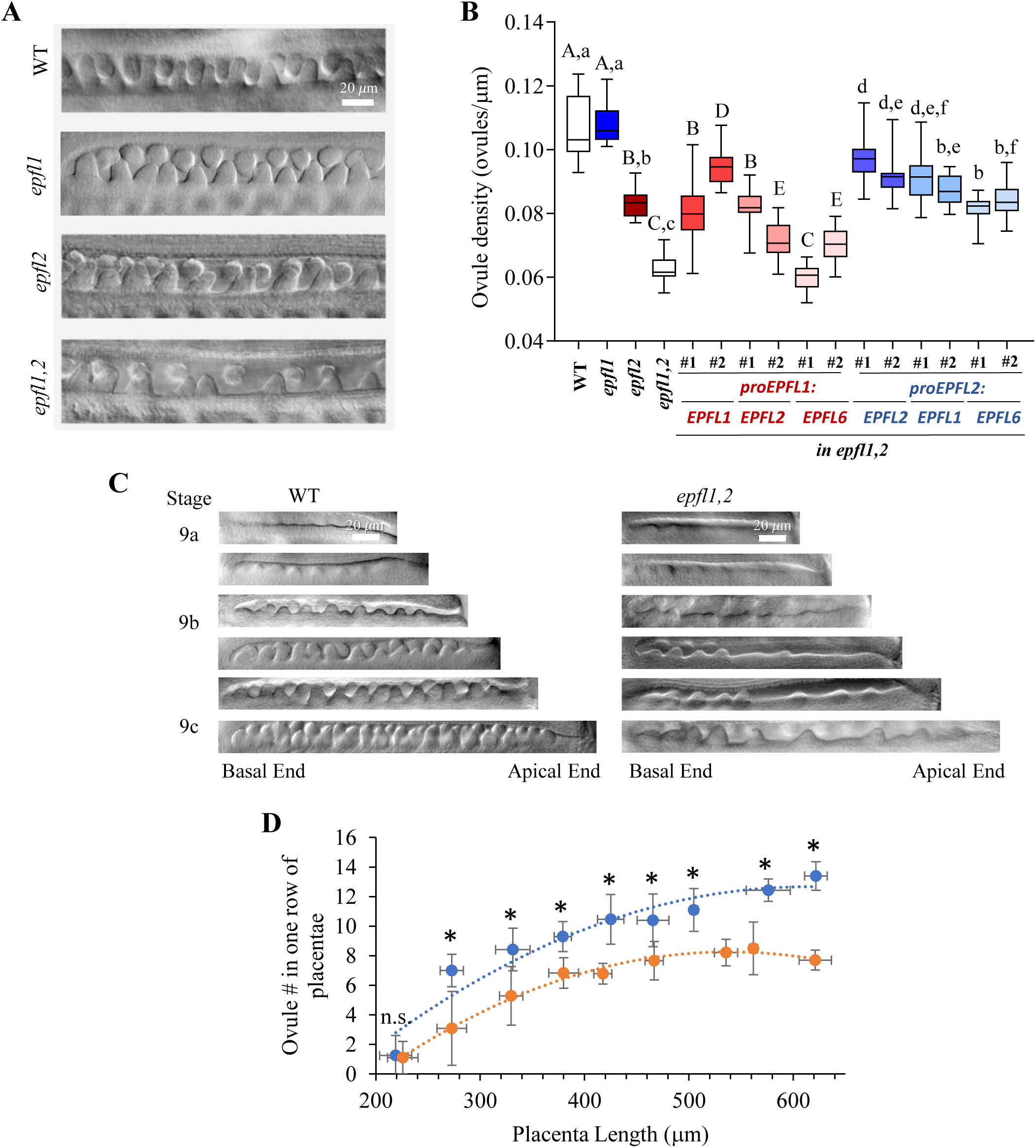
EPFL1 and EPFL2 positively regulate ovule density in a partially redundant manner. A. DIC images of stage 10 gynoecium demonstrate differences in ovule density. All images are to scale. B. Ovule density in WT, *epfl* mutants, and transgenic plants expressing the indicated constructs in *epfl1 epfl2* (labeled *epfl1,2*). Two independent transgenic lines (labeled #1 and #2) were analyzed for each construct. N=20. Data is represented using box–whisker plots: the whiskers represent maximum and minimum values, and the boxes represent the upper quartile, exclusive median, and lower quartile. Data were analyzed by one-way ANOVA and Tukey post-hoc test (P<0.01). Statistical differences between the EPFL1 promoter and its wild-type and mutant controls are coded with upper-case letters, likewise for the EPFL2 promoter using lowercase letters. C. DIC images of ovule initiation from stages 9a to 9c. One or two rows of ovules are visible in images. All images are to scale. D. Measurements of ovule number as a function of placenta length from stage 9a through 10. Measurements were separated into 50 mm placenta length bins from 200 to 650 mm. n-10-15. n.s.= no statistically significant difference. The trendline is polynomial. Values were compared using an unpaired two-tailed Student’s t-test p<0.05.

To test whether mutations in *EPFL1* and *EPFL2* are directly responsible for changes in ovule density, *proEPFL1:EPFL1* and *proEPFL2:EPFL2* were introduced into *epfl1 epfl2* and two independent transgenic lines were analyzed (Fig. 1B and Fig. S1). Both constructs contain terminator regions of the corresponding genes (Kosentka et al., 2019). The *proEPFL1:EPFL1* construct fully rescued the *epfl1* phenotype: the *epfl1 epfl2* plants expressing this construct were indistinguishable from *epfl2* plants. *proEPFL2:EPFL2* partially rescued the *epfl1 epfl2* mutant. Ovule number and density were slightly reduced in plants carrying the *proEPFL2:EPFL2* construct compared to either the wild type or *epfl1,* suggesting that some regulatory elements might be missing in our construct. To address whether *EPFL1*, *EPFL2*, and the more distantly related *EPFL6* are functionally redundant on the protein level, the promoter of *EPFL1* was used to drive the expression of EPFL2 and EPFL6, and the promoter of *EPFL2* was used to drive the expression of EPFL1 and EPFL6 in the *epfl1 epfl2* mutant. The coding sequences included introns. Indeed, expression of both *EPFL2* and *EPFL6* under the *EPFL1* promoter rescued ovule density and ovule number at least in some transgenic lines (Fig. 1B and Fig. S1). Likewise, *EPFL1* and *EPFL6* expressed under the *EPFL2* promoter enhanced ovule initiation similarly to *EPFL2* (Fig. 1B and Fig. S1).

In Arabidopsis, ovule initiation occurs asynchronously, with new primordia arising between previously formed ones as the placenta elongates (Yu et al., 2020). To investigate whether *EPFL1* and *EPFL2* regulate the initiation of early or late primordia, ovule density was measured throughout floral stage 9 (Fig. 1C). Overall, we observed stronger defects in ovule initiation in the apical half of the placenta (Fig. 1C). Quantification of ovule number as a function of placenta length demonstrated that ovule initiation in *epfl1 epfl2* is significantly reduced starting from the first wave of ovules (Fig. 1D). This suggests that the mutant has fewer ovules because the placenta is less able to form them. In summary, EPFL1 and EPFL2 synergistically promote the placenta’s ability to form ovules. EPFL1, EPFL2, and EPFL6 can substitute for each other when expressed under appropriate cis-regulatory elements.

### EPFL1 and EPFL2 promote ovule outgrowth and funiculus elongation

Ovule development in Arabidopsis has been described previously (Schneitz et al., 1995; Vijayan et al., 2021). Ovules initiate as finger-like protrusions. Later, inner and outer integuments develop, and three distinct regions form along a proximal-distal axis: the funiculus, the chalaza, and the nucellus. We noticed that *epfl1 epfl2* ovules grew more slowly than the wild-type ovules. To obtain quantitative data, the height of ovules was measured during meiosis of the megaspore mother cell (MMC), when callose is deposited into the cell wall between megaspores. Callose was stained with the fluorescent dye aniline blue. We observed that the length of *epfl1 epfl2* ovules was only about 63% of the wild type during megasporogenesis (Fig. 2A and B). This change in length was primarily due to an extremely short funiculus in the mutant. The *epfl1* single mutant formed ovules of regular size, while *epfl2* formed ovules shorter than the wild type but longer compared to *epfl1 epfl2*. These results suggest that EPFL1 and EPFL2 regulate ovule growth synergistically, with EPFL2 playing a more prominent role.

**Figure 2.**
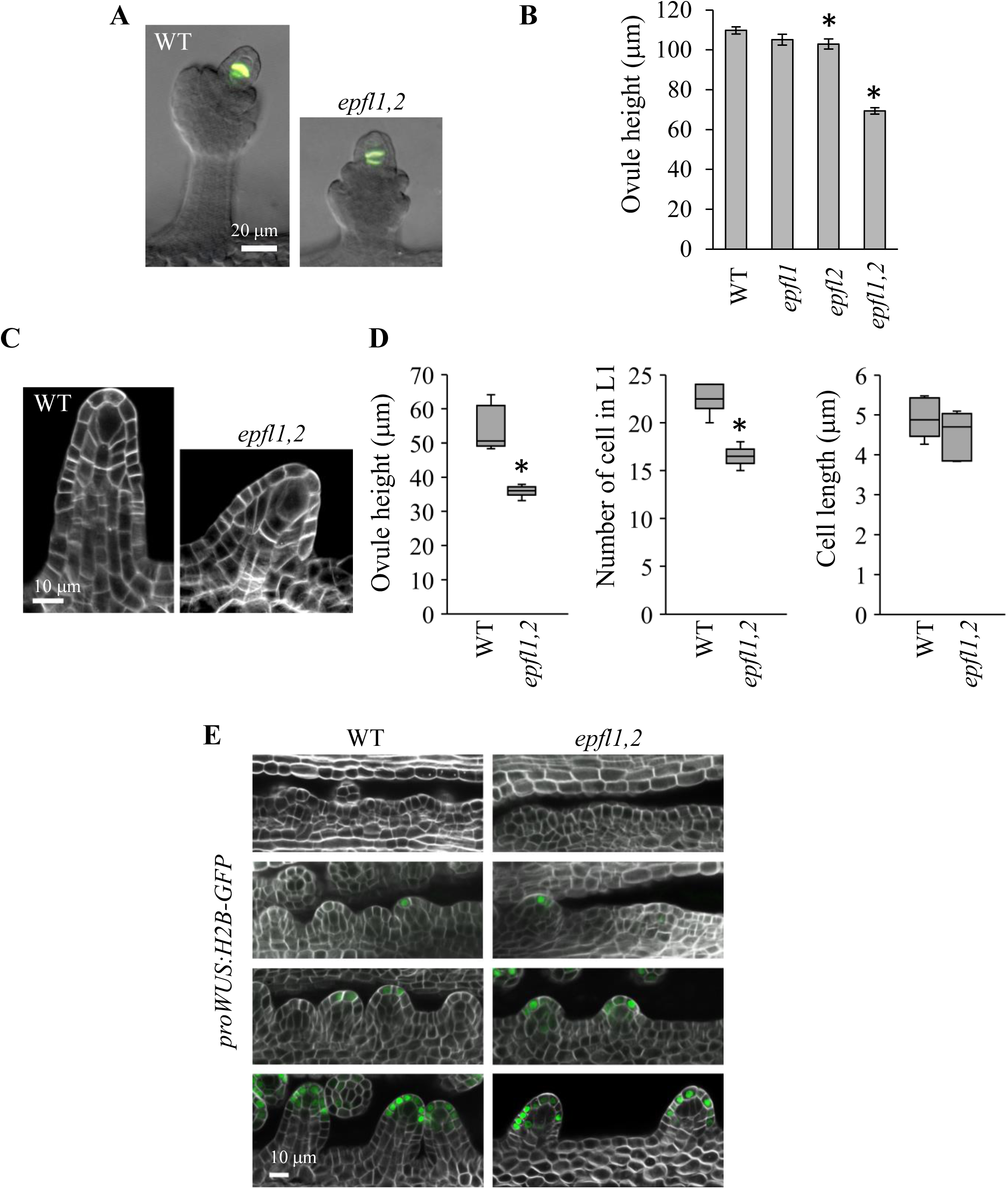
EPFL1 and EPFL2 promote early ovule growth. A. Epifluorescence and DIC overlay images of aniline blue stained ovules undergoing meiosis. B The mean height of ovules undergoing meiosis. n=10. Error bars=SE C. Representative confocal images of stage 2-I ovules with a characteristic enlarged megaspore mother cell. The cell walls were stained with SR2200. D. Ovule height (n=6), the number of L1 cells (n=6), and the length of L1 cells (15-24 cells/ovule in 6 ovules) were measured using cross-sections of six stage 2-I ovules. Data is represented using box–whisker plots: the whiskers represent maximum and minimum values, and the boxes represent the upper quartile, exclusive median, and lower quartile. B and D. Values that are significantly different from wt based on an unpaired two-tailed Student’s t-test (P<0.01) are indicated by asterisks. E. Representative confocal images of *proWUS:H2B-GFP* expression in ovules from stages 9a to 10. The cell walls were stained with SR2200. All images are under the same magnification in individual panels A, C, and E.

To quantify changes in ovule growth at an earlier stage, confocal cross sections of wild-type and *epfl1 epfl2* ovules were obtained at stage 2-I when a characteristic enlarged MMC forms (Fig. 2C). Already at this stage, *epfl1 epfl2* ovule height was about 67% of the wild type (Fig. 2D). We quantified number and length of epidermal layer (L1) cells using 2-D images of ovules. There was no statistically significant difference in the cell length, but the number of cells in the mutant was about 73% of the wild type (Fig. 2D). In the wild type, the transcription factor WUSCHEL (WUS) is expressed in the L1 layer at the top of the ovule starting from stage 1-II (Yu et al., 2020; Vijayan et al., 2021). Analysis of *proWUS:H2B-GFP* expression showed that *epfl1 epfl2* ovules were smaller than wild-type at stage 1-II when *WUS* expression consistently appeared. (Fig 2E, the third row from the top). At stage 2-I, *proWUS:H2B-GFP* demarcates the growing nucellus. Comparison of the wild type and *epfl1 epfl2* ovules at this stage suggested that while the nucellus was developing in the mutant, the funiculus was severely reduced (Fig 2E, the fourth row from the top). Overall, these results indicate that EPFL1 and EPFL2 regulate ovule outgrowth by promoting cell proliferation at its base, and they are critical for funiculus elongation.

### *EPFL1* and *EPFL2* show cell-type specific expression during ovule initiation

To understand the role of ERf/EPFL signaling during ovule development, we asked which cells produce ligands. We observed that *EPFL1* is expressed broadly in the placenta before ovule initiation (Fig. 3A), but with the onset of initiation, it is repressed in the subepidermal cell layer (Fig. 3B). Later, *EPFL1* expression is reduced at the tip of the ovule and in the boundaries (Fig. 3C and E). During integument initiation, *EPFL1* expression becomes further restricted to the basal side of the ovule (Fig. 3D). Previously, *EPFL2* expression was detected in the boundaries after ovules were initiated (Kawamoto et al., 2020). Our analysis indicated that *EPFL2* is very weakly expressed throughout the placenta before ovule initiation (Fig. 3F). During initiation, EPFL2 is upregulated, and its expression is restricted at first to boundaries between initiating ovules (Fig. 3G) and later spreads into the base of ovules (Fig. 3H). *EPFL2* expression does not surround the ovule at its base as *EPFL1* does but appears on the apical side of each ovule (Fig. 3I and J). In summary, our data suggest that while *EPFL1* and *EPFL2* have unique expression patterns, their expression overlaps at the base of the forming ovules.

**Figure 3.**
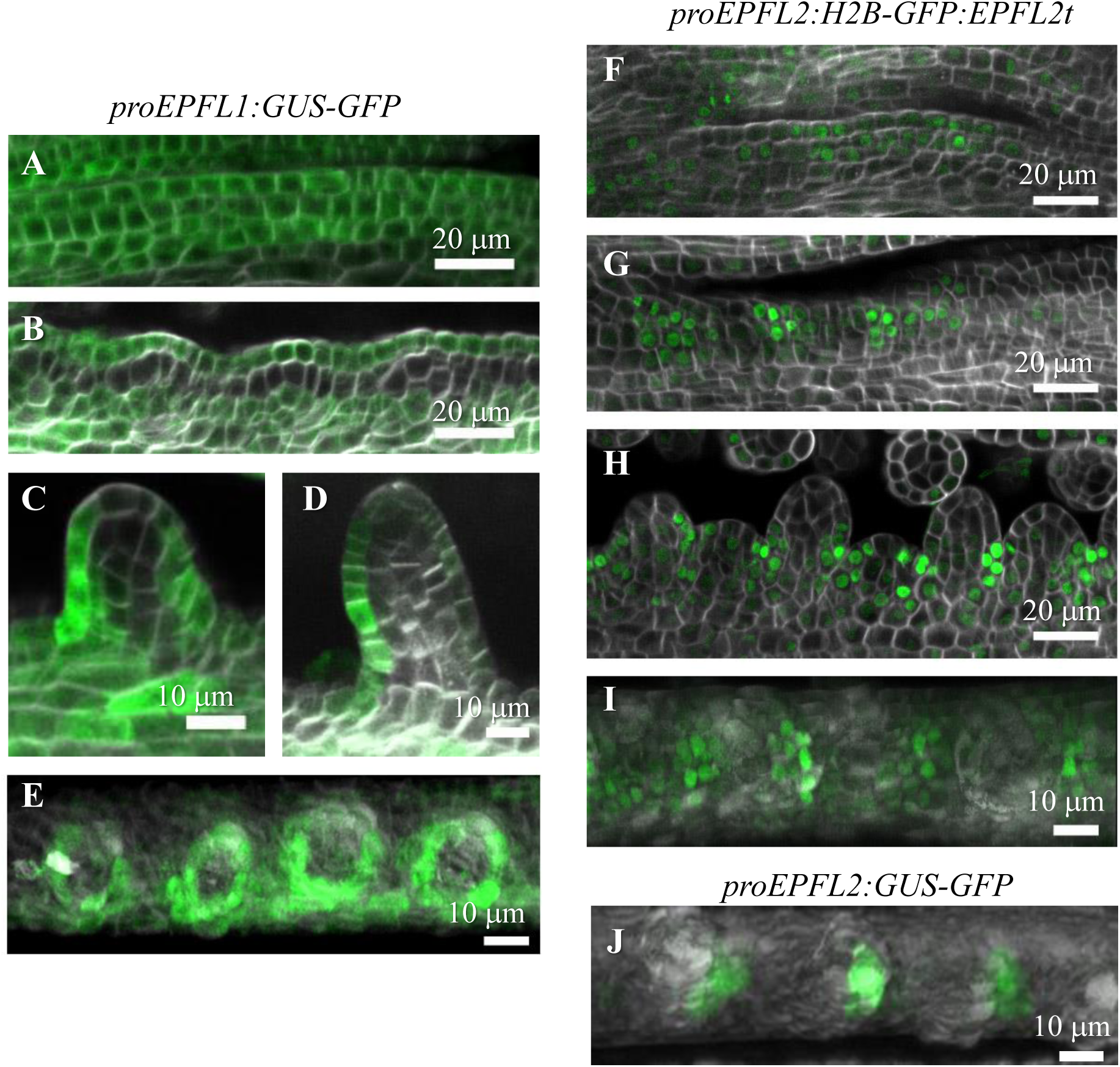
Expression patterns of *EPFL1* and *EPFL2* during early ovule development. Confocal images of the placenta at different stages of development in plants transformed with either *proEPFL1:GUS-GFP, proEPFL2:H2B-GFP: EPFL2term,* or *proEPFL2:GUS-GFP* (green). The cell walls were stained with SR2200 (white). A-D and F-H. Cross sections of *EPFL1* and *EPFL2* expression in ovules of different stages viewed from the side. E, I, and J. Confocal z-projections were used to show the expression of *EPFL1* and *EPFL2* in ovules from the top view.

### All three ER family receptors promote ovule initiation

Previous research studied the role of ERf receptors in the regulation of seed density using mature siliques (Kawamoto et al., 2020). In this stage, seed density depends on the spacing of ovules during initiation and the subsequent elongation of the carpel valves. To gain an insight into the role of receptors in ovule initiation, we analyzed flowers at stage 10, shortly after ovules were initiated. Measurements of ovule number and placenta length in *er* flowers showed a slight reduction in the placenta length but no changes in ovule density (Fig. 4A and B). Out of double *erf* mutants, only *erl1 erl2* had a small but statistically significant decrease in ovule density (Fig.4A). When all three receptors were disrupted, the ovule density was reduced to the *epfl1 epfl2* levels (Fig. 4A). This analysis suggests that all three receptors promote ovule initiation, with ERL1 and ERL2 playing slightly more prominent roles. ER is more important for the control of placenta elongation, but again, all three receptors control this process based on the extremely short placenta in the *er erl1 erl2* mutant (Fig. 4B).

**Figure 4.**
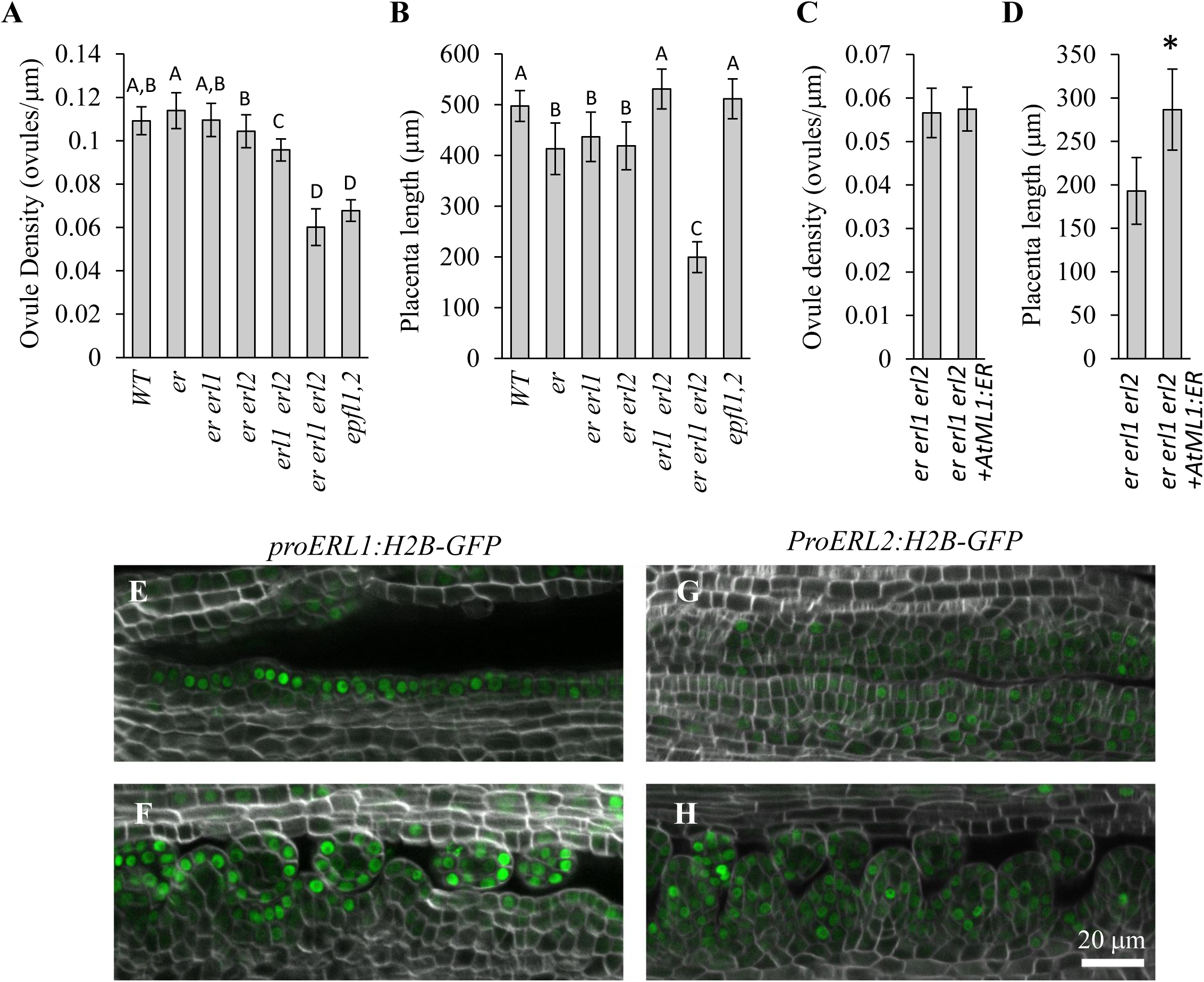
ERf receptors redundantly regulate ovule initiation. A-D. Mean ovule density and placenta length in stage 10 gynoecia of indicated genotypes. N=20. A one-way ANOVA using Tukey’s test with multiple comparisons was used to determine significance with a P-value < 0.01. **E-H** Representative confocal images of the placenta in *Landsberg erecta* plants transformed with either *proERL1:H2B-GFP* or *proERL2:H2B-GFP* (green). The cell walls were stained with SR2200 (white). All images are under the same magnification.

Previously, it was shown that *ER* and *ERL2* are expressed broadly in the placenta (Kawamoto et al., 2020). *ERL1* appeared only after ovules were formed, which is inconsistent with the observation that it plays a significant role in ovule initiation. We wanted to reexamine the expression of *ERL1*. For another project, we created transgenic lines expressing *proERL1:H2B-GFP* and *proERL2:H2B-GFP* in the Landsberg erecta ecotype. This ecotype contains a knockout mutation in *ER*. Here, we used these lines to test the expression of *ERL1* and *ERL2* in the absence of ER, which inhibits *ERL1* and *ERL2* expression (Pillitteri et al., 2007a). This experiment confirmed that *ERL2* is weakly and broadly expressed through the placenta (Fig. 4G and H). It also detected expression of *ERL1* in the L1 layer of the placenta during ovule initiation, supporting the role of ERL1 in that process (Fig. 4E and F). Because the *er erl2* mutant initiates ovules as efficiently as the wild type, we hypothesized that the expression of receptors in the L1 layer might be sufficient for ovule initiation. To test this hypothesis, we analyzed ovule initiation in *er erl1 erl2* plants that express *ER* under the epidermis-specific *AtML1* promoter (Sessions et al., 1999; Uchida et al., 2012). We observed that expression of ER in the epidermis can partially rescue the placenta elongation, but it has no effect on ovule density (Fig. 4C and D), which suggests that a broader expression of ERf receptors is necessary to promote ovule initiation. Maybe, in the *er erl2* mutant, *ERL1* is expressed outside of the epidermis. Another possibility is that a very small amount of ERL1 receptors in the internal layers, which we cannot detect by microscopy, is sufficient for the regulation of ovule initiation. Overall, the experiments reported so far indicate wide expression of all three receptors throughout the placenta.

### Decreased ovule initiation in *epfl1 epfl2* is not likely due to altered responses to auxin, cytokinin, or gibberellin

Multiple plant hormones have been implicated in the regulation of ovule number. Auxin is essential for ovule initiation (Benková et al., 2003; Bencivenga et al., 2012; Cucinotta et al., 2021), with auxin maxima being established at the tip of the ovule early in its formation (Galbiati et al., 2013). In Arabidopsis, cytokinins increase ovule number while gibberellins decrease it (Galbiati et al., 2013; Cucinotta et al., 2016; Gomez et al., 2018; Barro-Trastoy et al., 2020). To understand the cause of low ovule number in the *epfl1 epfl2* mutant, we tested whether its hormone metabolism is altered.

Analysis of *pDR5rev::GFP* expression suggested that auxin is present in ovules starting from flower stage 9c, ovule stage 1-II (Galbiati et al., 2013). Here, we used a more sensitive auxin reporter, *proDR5v2:ntdTomato*, that has a (TGTCGG)_9_ binding site with a higher affinity for ARFs compared to the (TGTCTC)_9_ binding site in *DR5rev* (Liao et al., 2015). Analysis of *proDR5v2:ntdTomato* expression indicated that discrete auxin maxima appear in the epidermis during stage 9a when ovules are barely noticeable (Fig. 5A). Auxin maxima first appear at the basal end of the placenta, consistent with the primacy of ovule initiation at that end. In the wild type, auxin accumulates in two to three cells at the tip of the ovule. The *epfl1 epfl2* mutant also accumulates auxin at the tip of the ovule, but the peaks are broader (Fig. 5A, compare the bottom row of the wild type and the mutant). This result is consistent with our previous work that demonstrated the negative effect of ERf signaling on auxin accumulation in the L1 layer of the SAM (Chen et al., 2013; DeGennaro et al., 2022). Next, we used the *proRPS5A:DII-n3Venus/proRPS5A:mDII-ntdTomato* reporter that is independent of signaling downstream of auxin receptors (Liao et al., 2015). Because auxin causes the proteolytic turnover of any protein harboring the DII degron sequence, this reporter indicates auxin accumulation through an absence of n3Venus fluorescence. Cells in which the reporter is expressed are marked by red ntdTomato fluorescence expressed from the *RPS5A* promoter (Fig. S3). This experiment confirmed that auxin forms a peak at the top of an ovule in both the wild type and the *epfl1 epfl2* mutant (Fig. S3).

**Figure 5.**
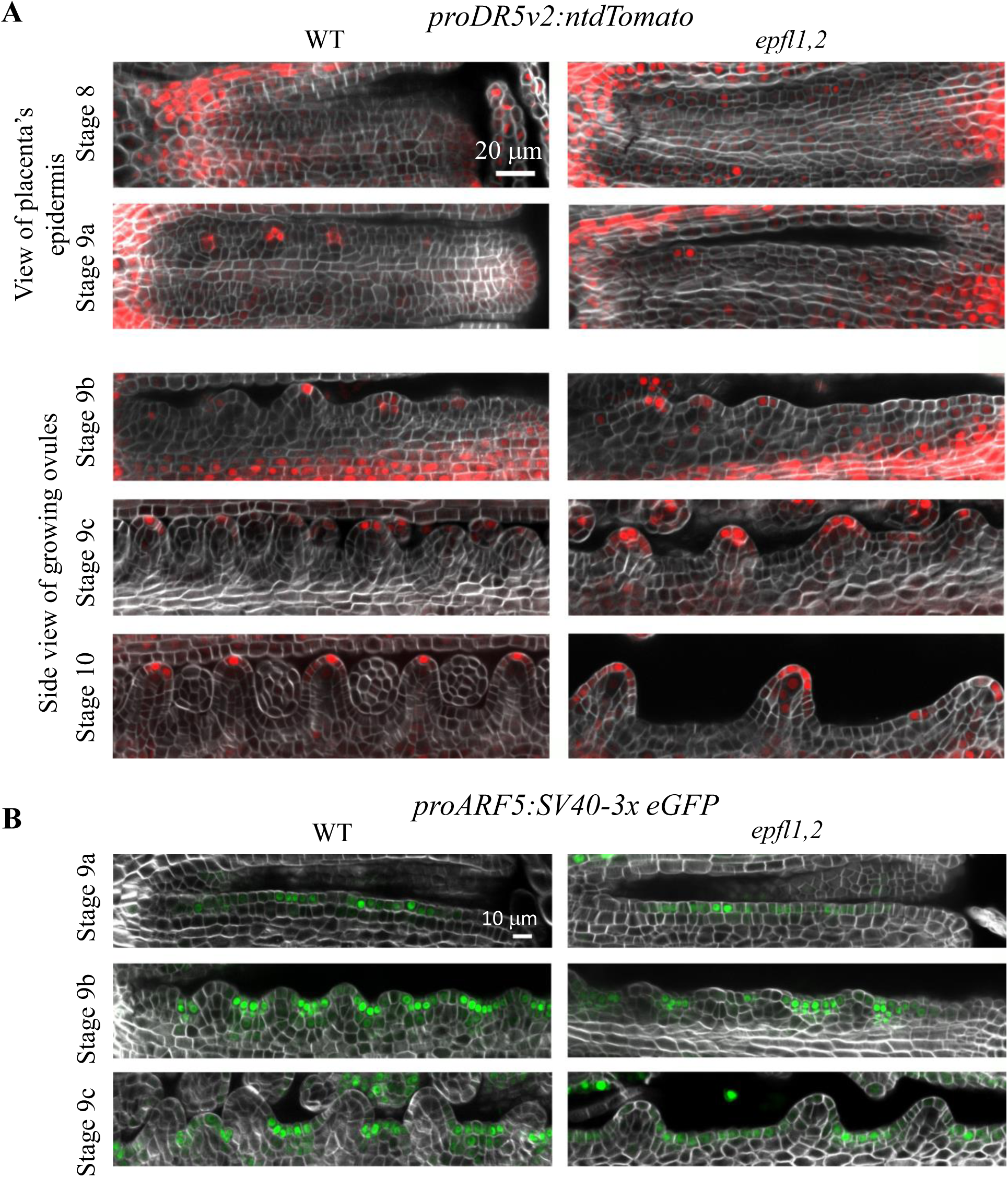
Auxin distribution and responses are similar in the wild type and *epfl1 epfl2*. A.. Representative confocal images of the placenta at different stages of development in plants transformed with *proDR5v2:ntdTomato* (red) indicated that auxin accumulates into maxima during ovule initiation in the *epfl1 epfl2* mutant (B) Representative confocal images of the placenta at different stages of development in plants transformed with proARF5:SV40-3x eGFP (green) indicated that transcription factor ARF5 is expressed similarly in the wild type and *epfl1 epfl2*. SV40 is a nuclear localization signal. A and B The cell walls were stained with SR2200 (white). All images in a panel are under the same magnification.

PIN1 is an auxin efflux carrier that is important for formation of auxin maxima in ovules (Okada et al., 1991; Benková et al., 2003; Bencivenga et al., 2012). As described previously, it is expressed in the placenta epidermis before ovule initiation (Galbiati et al., 2013). During ovule initiation, its expression in the epidermis is upregulated at the center of the ovule and decreased in the boundaries (Fig. S4A). PIN1 expression is absent in the subepidermal layer of the placenta but is present in the deeper layers (Fig. S4B). Over time, PIN1 expression in the deeper layers is restricted to the forming vasculature (Fig. S4C). PIN1 is polarly localized in the plasma membrane to enable auxin transport to the ovule tip (Fig. S4D). A comparison of PIN1 expression in the wild type and the *epfl1 epfl2* mutant did not identify any differences in its expression pattern or polar localization (Fig. S4).

Previously, treatment of *er erl1 erl2* seedlings with N-1-naphthylphthalamic acid (NPA), an inhibitor of polar auxin transport, or with synthetic auxin 2,4-Dichlorophenoxyacetic acid (2,4D) partially rescued leaf initiation because, in both cases, it increased accumulation of auxin in the L1 layer (DeGennaro et al., 2022). Here, a treatment of flowers with 10 μM NPA reduced ovule initiation in both the wild type and the *epfl1 epfl2* mutant (Fig. S5A), while 50 μM 2,4 D did not alter ovule initiation in either genotype (Fig. S5B). Altered gravitropism of internodes in response to 2,4D treatment served as a positive control, confirming that a physiologically relevant concentration of hormone was used. This result highlights differences in auxin homeostasis between the shoot apical meristem and the placenta, but most importantly, it indicates the similarity of auxin responses in the wild type and the *epfl1 epfl2* mutant.

In the presence of high auxin, AUX//IAA proteins are degraded, relieving repression of transcription factors such as MP/ARF5. Previous research showed that *pMP:MP-GFP* is expressed throughout the placenta before ovule initiation (Galbiati et al., 2013). After ovules initiate, MP/ARF5 expression is suppressed at the tip of the ovule, and it is expressed in the boundaries. Our analysis of *proARF5:SV40-3xeGFP* (Rademacher et al., 2011) expression indicated that starting from early stage 9a, MP/ARF5 is being restricted to boundaries of ovules and is expressed similarly in the wild type and the *epfl1 epfl2* mutant (Fig. 5B). Altogether, these data point towards similar auxin distributions and responses during ovule initiation in the wild type and the *epfl1 epfl2* mutant.

Previously, the exogenous application of the synthetic cytokinin 6-benzylaminopurine (BAP) led to increased placenta length and ovule number in wild-type plants (Galbiati et al., 2013; Cucinotta et al., 2016). Our treatment of the wild type and *epfl1 epfl2* inflorescences with 1mM BAP led to a significant increase in ovule number and placenta length without significant changes in ovule density (Fig. S6). This suggests that exogenous cytokinins promote the elongation of the placenta, allowing initiation of more ovules, but do not regulate the spacing of ovule primordia along its length. The *epfl1 epfl2* mutant is responsive to cytokinin in a similar manner to the wild type, and the ovule density phenotype of the *epfl1 epfl2* mutant is not restored after BAP treatment (Fig. S6). This suggests that the *epfl1 epfl2* placenta does not have an impaired response to cytokinin.

Gibberellins (GA) inhibit ovule initiation (Gomez et al., 2018; Barro-Trastoy et al., 2020). As expected, the application of GA to wild-type inflorescences caused a decrease in ovule number and density without significantly affecting placenta length, while the application of the gibberellin biosynthesis inhibitor paclobutrazol (PAC) caused increased ovule number and density without altering placenta length (Fig. S6). The wild-type and *epfl1 epfl2* responses to GA and PAC were similar (Fig. S6), suggesting that changes in ovule initiation in the mutant are not caused by altered GA signaling or biosynthesis.

### Synergistic regulation of ovule initiation by *CUC2, CUC3*, *EPFL1,* and *EPFL2*

CUP-SHAPED COTYLEDON (CUC) transcription factors regulate ovule spacing (Galbiati et al., 2013; Gonçalves et al., 2015). We examined genetic interactions between *EPFL1, EPFL2, CUC2,* and *CUC3 b*ecause these two CUC proteins are expressed in the boundaries of initiating ovules in a pattern similar to *EPFL2* (Gonçalves et al., 2015). Relative to the wild type, the ovule density was not changed in single *cuc2* or *cuc3* mutants and was slightly increased in the *cuc2 cuc3* double mutant (Fig. 6A). The addition of *cuc3* to the *epfl1 epfl2* mutant significantly reduced ovule density relative to *epfl1 epfl2* (Fig. 6A and B), indicating a synergistic interaction between *CUC3*, *EPFL1* and *EPFL2* and the importance of *CUC3* for ovule initiation. The addition of the *cuc2* mutation to either *epfl1 epfl2* or *epfl1 epfl2 cuc3* did not alter ovule density of those mutants (Fig. 6A). At the same time, we noticed that ovules in the *epfl1 epfl2 cuc2 cuc3* mutant looked dramatically different. There was a delay in ovules outgrew from the placenta compared to the wild type or lower-order mutants (Fig. 6B and C). In addition, during megasporogenesis, ovules were broad and short without a clearly defined funiculus (Fig. 6D). This indicates that *CUC2* and *CUC3* regulate the morphology of ovules in synergy with *EPFL1* and *EPFL2* and promote the elongation of the funiculus. *EPFL1, EPFL2, CUC2*, and *CUC3* also redundantly promote the elongation of the placenta (Fig. S7).

**Figure 6.**
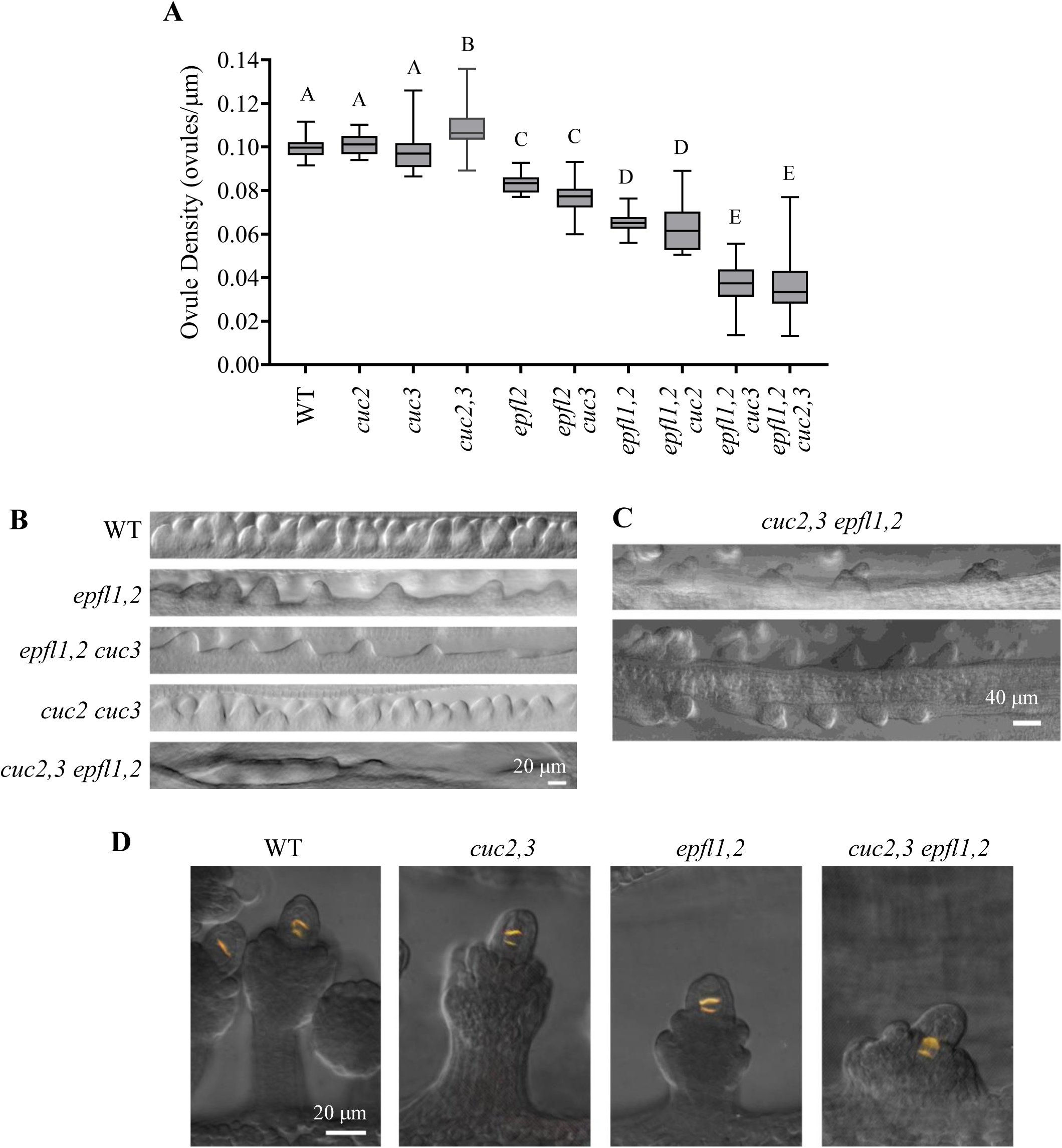
EPFL1,2 and CUC2,3 regulate ovule initiation and growth synergistically. A. Ovule density measurements of the wild type and indicated mutants. N=10. Data is represented using box– whisker plots: the whiskers represent maximum and minimum values, and the boxes represent the upper quartile, exclusive median, and lower quartile. A one-way ANOVA using Tukey’s test with multiple comparisons was used to determine significance with a P-value < 0.01. B. DIC images of stage 10 gynoecia demonstrate differences in ovule initiation in indicated genotypes. C. DIC images of stage 11 gynoecia indicate that ovules can initiate in the *cuc2,3 epfl1,2* mutant over time. D. Epifluorescence and DIC overlay images of aniline blue stained ovules undergoing meiosis demonstrate altered morphology of ovules in indicated mutants. B-D Images in the same panel are under the same magnification.

We tested whether EPFLs and CUCs influence each other’s expression and found that the expression of *CUC3* was expanded in the placenta of the *epfl1 epfl2* mutant. *EPFL2* and *CUC2* are expressed in a similar pattern. They are present between ovules and at the base of ovules and are excluded from the forming nucellus (Fig. S8). It has been reported previously that CUC2 inhibits *EPFL2* expression (Li et al., 2020), but we did not observe significant changes in the expression of *EPFL2* in the *cuc2 cuc3* mutant (Fig. S8). We did not detect changes in *CUC2* expression in the *epfl1 epfl2* mutant (Fig. S8). In the wild type, *CUC3* expression is efficiently restricted to the boundaries between forming ovules (Fig. 7, Fig. S9). In the *epfl1 epfl2* mutant, *CUC3* was expressed more broadly, which is partly due to broader boundaries between ovules in the mutant (Fig. S9, stages 9c and 10). In addition, we observed the reduced inhibition of *CUC3* expression in the nascent ovules of the mutant (Fig. 7A, stages 9a and 9b). This was specific to *CUC3*, as we did not observe a delay in the repression of another boundary transcription factor, *MP/ARF5* (Fig. 5B).

**Figure 7.**
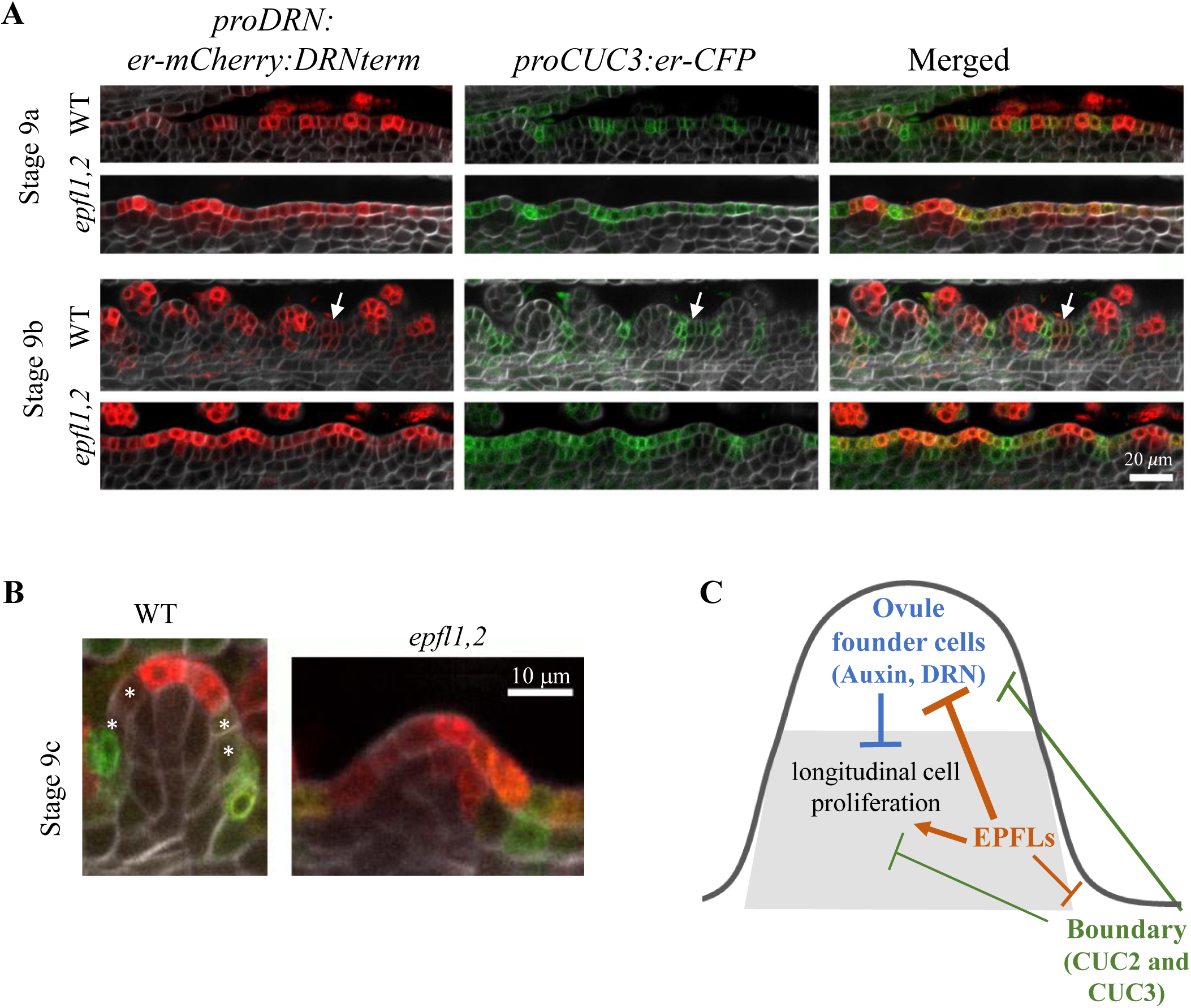
EPFL1 and EPFL2 are restricting the expression of DRN and CUC3. A. During ovule initiation, the expression of *DRN* and *CUC3* overlaps in more cells and for a longer period in the *epfl1 epfl2* mutant compared to the wild type. B. Cells without *DRN* and *CUC3* form close to the base of the ovule at stage 9c (indicated by asterisk) in the wild type but not in *epfl1 epfl2*. A and B. Representative confocal images of the wild type and *epfl1 epfl2* placenta at different stages of development in plants co-transformed with *proDRN:er-mCherry:DRNterm* (red) and *proCUC3:er-CFP* (green). The cell walls were stained with SR2200 (white). All gynoecia are oriented with a basal end on the left and an apical end on the right. A white arrow indicates the upregulation of *DRN* and CUC3 expression in a few boundary cells that will become an ovule. All images in this figure are under the same magnification. C. Model of EPFL function during ovule initiation. During placenta patterning, cells differentiate into ovule founder cells and boundary cells. In both of these areas, longitudinal cell proliferation is slow. EPFLs inhibit both ovule founder cell and boundary cell identities, allowing the formation of an area of longitudinal cell proliferation and efficient ovule bulging from the placenta. CUC2 and CUC3 are also able to inhibit ovule founder cell identity.

The transcription factor BZR1 controls organ boundary formation by directly repressing *CUC* genes (Gendron et al., 2012). A dominant *bzr1-1D* mutation increases BZR1 stability (Wang et al., 2002) and, thus, should downregulate *CUC* expression. To understand the role of BZR1 in ovule initiation, we investigated genetic interactions between *epfl1 epfl2* and *bzr1-1D*. We observed that the *bzr1-1D* mutation decreases ovule initiation in the *epfl1 epfl2* mutant but not in the wild type (Fig. S10) similar to *cuc3* (Fig. 6A). We speculate that the repression of *CUC3* by *bzr1-1D* is the cause of the similarity of *bzr1-1D epfl1 eplf2* and *cuc3 epfl1 epfl2* phenotypes. In summary, our data suggest that *EPFL* and *CUC* genes have common targets in the placenta and that BZR1 might negatively regulate boundary formation during ovule initiation.

### EPFL1 and EPFL2 stabilize the patterning of the placenta

The transcription factor *DRN* is the earliest marker of ovule initiation. At first, it is expressed throughout the L1 layer of the placenta, but then it is efficiently suppressed in the ovule boundaries during the earliest steps of ovule initiation (Kirch et al., 2003; Kawamoto et al., 2020). In the *epfl2* mutant, the spacing of *DRN* expression in the placenta during ovule initiation is variable, indicating that EPFL2 patterns the placenta by suppressing *DRN* between incipient ovules (Kawamoto et al., 2020). To investigate *DRN* expression in the *epfl1 epfl2* mutant, we first analyzed the expression of the *proDRN:H2B-GFP:DRNterm* reporter. In the wild type, *DRN* is very efficiently suppressed in the boundaries between ovules during early stage 9a before ovule primordia protrude from the placenta (Fig. S11). During wild-type ovule outgrowth, *DRN* expression is restricted to only a few L1 cells on top of a primordium (Fig. S11). The formation of new ovules between existing ones is also marked by *DRN* expression in boundary regions. In the *epfl1 epfl2* mutant during stages 9a and 9b, *DRN* expression is not suppressed in the boundary regions and, instead, remains broadly distributed in the L1 cells of the placenta even after the protrusion of ovule primordia (Fig. S11). Only, by stage 10, right before integuments are ready to form, is *DRN* expression in *epfl1 epfl2* finally restricted to the top of the primordia.

Next, we analyzed the expression of *DRN* using the *proDRN:er-mCherry:DRNterm* reporter (Fig. 7A), which allowed us to image the expression of *DRN* and *CUC3* simultaneously. In the wild type, the expression of these two genes overlapped in a few cells during stage 9a on the apical side of a forming ovule. Once the ovule visibly protruded from the placenta in stage 9b, there was no overlap of *DRN* and *CUC3* expression except in the boundary regions where a new ovule would form. In the *epfl1 epfl2* mutant, *DRN* and *CUC3* were co-expressed not only during the 9a stage but also during 9b. During the 9c and 10 stages, *CUC3* became restricted to boundaries, but *DRN* was still broadly expressed in boundaries and most L1 cells of protruding ovules, including the growing funiculus (Fig. 7B and Fig. S9). In summary, during ovule initiation, the wild-type L1 placenta cells acquire different identities, with *DRN* expression becoming restricted to the top of the ovule and *CUC3* to the few cells between ovules. In a clearly protruding ovule, the majority of L1 cells, excluding the top cells, do not express either of the markers. This pattern is changed in the *epfl1 epfl2* mutant, which suggests that EPFL1 and EPFL2 promote the separation of *CUC3* and *DRN* expression and the formation of a new cell type at the base of the ovule. Thus, EPFL/ERf signaling enables cell-to-cell communication in the placenta and is responsible for the establishment of unique cell identities.

## Discussion

While ERf receptors redundantly regulate many different aspects of plant development (Shpak, 2013; Jiang et al., 2022), their EPF/EPFL ligands have unique functions. For example, stomata development is regulated by EPF1, EPF2, and STOMAGEN/EPFL9 (Hara et al., 2007; Hara et al., 2009; Hunt and Gray, 2009; Hunt et al., 2010; Sugano et al., 2010), the shoot apical meristem structure, leaf initiation, and plant height are regulated by EPFL1, EPFL2, EPFL4, and EPFL6 (Abrash et al., 2011; Uchida et al., 2012; Kosentka et al., 2019), and early cotyledon growth and development of leaf teeth are regulated by EPFL2 (Tameshige et al., 2016; Fujihara et al., 2021).

Here, we studied the role of the ERf signaling pathway in ovule initiation, including control of placenta length and ovule spacing. The expression data indicate that ERf signaling functions broadly in the placenta before ovules initiate, but after initiation, its function is limited to the spaces between ovules and the elongating ovule base. All three ERf receptors and two ligands, EPFL1 and EPFL2, regulate ovule spacing. Placenta elongation is controlled by all of the receptors and by at least four ligands: EPFL1, EPFL2, EPFL4, and EPFL6. As the placenta is slightly longer in *epfl1 epfl2 epfl4 epfl6* compared to *er erl1erl2*, either additional ligands regulate placenta elongation or receptors can regulate it in a ligand-independent manner. While the three receptors function synergistically, ERL1 and ERL2 are more crucial for ovule spacing, whereas ER plays a more significant role in placenta elongation. Our data do not contradict a previously published report that ERL1 and ERL2 function antagonistically to ER in regulating seed density (Kawamoto et al., 2020). While all three receptors increase ovule density during the ovule initiation step, ER-induced elongation of siliques later in development promotes reduced seed density.

We also observed that EPFL1 and EPFL2 promote the elongation of nascent ovules by enhancing cell proliferation at their base. This fits into a common pattern as the role of ERf signaling in the regulation of cell proliferation is well-known. ERf signaling promotes the proliferation of cells along the apical-basal axis in internodes, pedicels, floral organs, leaves, and cotyledons (Shpak et al., 2003; Shpak et al., 2004; Abraham et al., 2013; Patel et al., 2013; Chen and Shpak, 2014). A detailed analysis of pedicel growth revealed that ERf signaling accelerates the cell cycle by shortening the time required for a cell to double in size (Bundy et al., 2012). The EPFL1/ EPFL2-induced proliferation of cells at the base of an ovule leads to a longer funiculus. Few mutants with a shorter funiculus have been described. To our best knowledge, shorter funiculi were previously observed only in higher-order *epfl* mutants, the *er erl1 erl2* mutant, the *serk1 serk2 serk3* mutant (Li et al., 2023), and the *cuc1 cuc2* mutant (Ishida et al., 2000), all genes with a close relationship to the ERf pathway. Thus, ERf signaling might be the main regulator of funiculus length. The role of ERf signaling in the elongation of the ovule’s funiculus is analogous to its role in the elongation of the leaf’s petiole and the flower’s pedicel; in each case, cell proliferation is induced at the base of the forming organ, which leads to the formation of a stalk-like structure.

Ovule initiation is controlled by auxin, cytokinin, and gibberellins (Yu et al., 2022). While ERf signaling is associated with all three hormonal pathways in a variety of developmental processes (Woodward et al., 2005; Chen et al., 2013; Uchida et al., 2013; Tameshige et al., 2016; DeGennaro et al., 2022; Zhao et al., 2023), we did not find a connection during ovule initiation. Throughout our studies, we noted a distinct role for each hormone in ovule initiation. Cytokinins are crucial for placenta elongation, whereas gibberellins and auxin control ovule spacing in opposing ways. Previously, the earliest auxin maxima were detected only at stage 1-II of ovule development in finger-like protrusions (Ceccato et al., 2013). Using more sensitive *proDR5v2:ntdTomato* and *proRPS5A:DII-n3Venus/proRPS5A:mDII-ntdTomato* reporters, we were able for the first time to detect the formation of auxin maxima in the placenta right before ovule initiation which supports auxin’s role in specifying ovule founder cells.

A recurring theme in ERf signaling is its synergy with boundary genes. Previously, we demonstrated that ERf receptors regulate cotyledon and leaf initiation synergistically with the kinase PID, which is expressed in the boundaries and is important for organ separation (Furutani et al., 2004; Landrein et al., 2015; DeGennaro et al., 2022). Here, we show the synergy of *EPFLs* with the major determinants of the boundary domain, the *CUC* genes. The *cuc3* mutation does not alter ovule initiation on its own but strongly decreases it in the *epfl1 epfl2* background. *EPFL1, EPFL2, CUC2*, and *CUC3* synergistically regulate funiculus elongation. The synergy of *bzr1-1d* with *epfl1 epfl2* can be also explained by the role of brassinosteroids in boundary formation (Gendron et al., 2012). While brassinosteroids promote ovule initiation (Huang et al., 2013), the upregulation of brassinosteroid signaling by *bzr1-1D* mutation in the *epfl1 epfl2* background led to its severe disruption. This outcome can be understood if brassinosteroids inhibit the expression of *CUC* genes in the placenta much like they do in the shoot apical meristem (Gendron et al., 2012).

*CLAVATA3 (CLV3)* and *WUS* are the main targets of EPFLs in the shoot apical meristem (Liu et al., 2020; Zhang et al., 2021; Uzair et al., 2024). However, neither *CLV3* nor *WUS* is expressed in the placenta during the specification of ovule initiation sites. EPFL2 inhibits auxin responses in the elongating sides of a leaf tooth (Tameshige et al., 2016), but broader auxin responses in *epfl1 epfl2* ovules were observed only after ovules had already been initiated. Moreover, our previous work on leaf initiation suggested that increased accumulation of auxin in *er erl1 erl2* mutants is not the cause of decreased organ primordia initiation (DeGennaro et al., 2022). To understand the role of ERf signaling in ovule initiation, we analyzed the earliest known markers of cell differentiation in the placenta: *DRN* has been previously identified as an early marker of ovule founder cells (Kawamoto et al., 2020) while *CUC3* is an early marker of boundary regions (Gonçalves et al., 2015). In the wild type, these two genes are expressed in different epidermal cells during the specification of ovule sites in the placenta. As soon as the ovule bulges out of the placenta, *DRN*- and *CUC3*-expressing cells are separated by cells that express neither. We observed that the separation of *DRN* and *CUC3* expression to different cells and the appearance of cells without these two markers is impaired in the *epfl1 epfl2* mutant.

Here, we propose a new model that is consistent with all known data to date. After the placenta reaches a specific length, its cells differentiate into ovule founder cells and boundary cells. As CUC transcription factors are central to ovule spacing (Galbiati et al., 2013; Gonçalves et al., 2015), they should play a major role in defining the size of boundaries and the inhibition of ovule founder cell identity. Based on its expression pattern, DRN is a good candidate for ovule founder fate induction. Once patterning is established, it is reinforced by EPFL1 and EPFL2, which strengthen the separation of the ovule founder and boundary cell identities and promote the formation of cells with a new identity in the middle (Fig. 7C). The emergence of these new cells is central to efficient ovule bulging because both ovule founder and boundary cells are characterized by slow cell proliferation. The synergy of *epfl* and *cuc* mutations suggests that, for efficient ovule initiation, inhibition of ovule founder cell identity by ERf signaling is more important than inhibition of boundary fate.

We previously observed that ERf signaling restricts *DRN* and *DRNL* expression in the shoot apical meristem (Uzair et al., 2024), and now we have determined that it limits *DRN* expression in the placenta. Is *DRN* the main target of EPFL1 and EPFL2 during ovule initiation? Unfortunately, we do not know the answer to this question, as ERf signaling might be targeting multiple genes to restrict ovule founder cell identity, and DRN’s role in ovule initiation has not yet been studied at the genetic level. It is also unclear how *DRN* expression is regulated during ovule initiation. At the tips of the cotyledons, auxin via MP/ARF5 promotes the expression of *DRN* (Cole et al., 2009). But in the shoot apical meristem *DRN* expression is inhibited by MP/ARF5 (Luo et al., 2018). During ovule initiation, *DRN* and auxin are expressed in the ovule founder cells, while *MP/ARF5* is expressed in the boundary. Interestingly, in the *epfl1 epfl2* placenta, the expression of *MP/ARF5* and the accumulation of auxin have not changed, but *DRN* expression has, suggesting an auxin-independent role for EPFLs in the regulation of *DRN* expression.

The initiation of all aboveground organ primordia starts with the specification of organ founder cells and the delineation of boundary regions followed by organ growth along the apical-basal axis. ERf signaling regulates the elongation of all aboveground organs. The separation of organ founder and boundary cells with a cell proliferation zone might play a universal role in ERf signaling during organ initiation. This role would be similar to ERf function in the shoot apical meristem, where ERfs promote a fast-dividing peripheral cell identity positioned between slowly dividing cells of the central zone and the boundary. In the leaf epidermis, ERf signaling promotes cell proliferation by limiting the expression of the cell differentiation signal MUTE (Pillitteri et al., 2007b; Qi et al., 2017; Han et al., 2018; Zuch et al., 2023). Identifying the downstream targets of ERf signaling during early organ primordia growth could help us determine if the universal function of this signaling pathway is to inhibit transcription factors that induce cell fates with low proliferative potential.

## Materials and methods

### Plant Materials and Growth Conditions

All research was performed using the Columbia ecotype of *Arabidopsis thaliana*. The *epfl4 (chl-2)*, *epfl6 (chal-2), cuc2-3*, *cuc3-105, epfl1-1, epfl2-1,* and *bzr1-1D* mutants were described previously (Wang et al., 2002; Hibara et al., 2006; Abrash et al., 2011; Tameshige et al., 2016; Kosentka et al., 2019). The higher order mutants *epfl1 epfl2*, *epfl4 epfl6,* and *er-105 erl1-2 erl2-1* were generated earlier (Shpak et al., 2004; Kosentka et al., 2019). When the *cuc3-105* mutation was outcrossed into *epfl2-1* backgrounds, the *epfl2-1* mutation was genotyped according to (Kosentka et al., 2019). The *cuc2-3* and *cuc3-105* mutations were identified based on genotyping using three-primer PCR. For genotyping of *cuc2-3*, the primers CUC2 GT F (5’-GCACGCACGCATACTCAGATAGA-3’) and CUC2 GT R (5’-CCAGTAACCAGCCTCAGTTGCTC-3’) were used to amplify the 931bp wild type fragment, and CUC2 GT R and SAIL LB (5’-TAGCATCTGAATTTCATAACCAA-3’) were used to amplify an approximately 550 bp fragment of the mutant allele. For genotyping of *cuc3-105*, the primers CUC3 GT F (5’-GATGATGCTTGCGGTGGAAGATG-3’) and CUC3 GT R (5’-GACATCCACACCACACGTACAGG-3’) were used to amplify the 1062bp wild type fragment, and CUC3 GT R and GABI-Kat LB (5’-ATATTGACCATCATACTCATTGC-3’) were used to amplify an approximately 400 bp fragment of the mutant allele. Double *cuc2-3 cuc3-105* mutants were identified based on cotyledon fusions and confirmed by genotyping. During a cross with *epfl1-1 epfl2-1*, a homozygous *bzr1-1D* mutant was identified based on the increased size of leaves and flower organs, delayed flowering, and bending of the inflorescence stem. During crosses, the double *epf1-1 epfl2-1* mutant can be identified by its shorter petals and confirmed by genotyping (Kosentka et al., 2019).

Plants were grown on 1x Murashige and Skoog’s (MS) solid media supplemented with 1% (w/v) sucrose until 5-7 days post-germination when they were moved to the soil. The soil mixture contained a 1:1 ratio of Lambert LM-Org soil to Palmetto Fine-Medium A-2 vermiculite supplemented with 75 mg/100g of Miracle-Gro and 0.25 g/100g of Osmocoat 15-9-12 (Scotts). Plants were grown at 20^0^ C under long-day conditions of 18 hours of light and 6 hours of darkness.

### Generation of Transgenic Plants

Plasmids containing *proEPFL1:EPFL1:EPFL1term* (pPZK411) and *proEPFL2:EPFL2:EPFL2term* (pPZK412) in the binary vector pPZP222 and *proEPFL1:GUS-GFP:35sTer*m (pPZK401) and *proEPFL2:GUS-GFP:35sTerm* (pPZK402) in the binary vector pKGWFS7 were described previously (Kosentka et al., 2019). To generate *proEPFL1:EPFL2:EPFL1term* (pAMO101), overlapping PCR was used to fuse the 2.05 kb promoter of *EPFL1* with the transcribed region of *EPFL2* followed by the 0.95 kb *EPFL1* terminator. The fragment was inserted into pPZP222 between the BamHI and KpnI sites. To generate *proEPFL2:EPFL1:EPFL2term* (pAMO100), overlapping PCR was used to fuse the transcribed region of *EPFL1* with the 1.1 kb *EPFL2* terminator. The fragment was inserted into pPZP222 between the BamHI and KpnI sites. The 2.6 kb *EPFL2* promoter was then inserted between the HindIII and BamHI sites of the vector. To generate *proEPFL1:EPFL6:EPFL1term* (pAMO105), overlapping PCR was used to fuse the 2.05 kb promoter of *EPFL1* with the transcribed region of *EPFL6* followed by the 0.95 kb *EPFL1* terminator. The fragment was inserted into pPZP222 between the BamHI and KpnI sites. To generate *proEPFL2:EPFL6:EPFL2term* (pMIG101), overlapping PCR was used to fuse the transcribed region of *EPFL6* with the 1.1 kb *EPFL2* terminator. The fragment was inserted into pPZP222 between the BamHI and KpnI sites. The 2.6 kb *EPFL2* promoter was then inserted between the HindIII and BamHI sites of the vector. To generate *proEPFL2:H2B-eGFP:EPFL2term* (pAMO124), *H2B-eGFP* was fused with the 1.1 kb *EPFL2* terminator using overlapping PCR. This fragment was inserted into pPZP222 between the BamHI and KpnI sites. Then the 2.6 kb *EPFL2* promoter was inserted into the plasmid between the HindIII and BamHI sites. The template for amplifying the H2B-EGFP sequence was a plasmid from the Nimchuk lab (UNC Chapel Hill, USA). To generate the *proDRN:H2B-eGFP:DRNterm* construct (pAMO102b), *H2B-eGFP* was fused to the 1.38 kb *DRN* terminator using overlapping PCR and inserted into the binary vector pPZP222 between the BamHI and SalI sites. Then, the 4.86 kb *DRN* promoter was inserted using the BamHI site. To generate *proDRN:er-mCherry:DRNterm* (pAMO119), the 4.86 kb *DRN* promoter was inserted into pPZP222 between the BamHI and SalI sites. Next, *er(endoplasmic reticulum)-mCherry* was fused with the 1.38 kb *DRN* terminator using overlapping PCR and inserted into the plasmid using SalI. The *er-mCherry* sequence was amplified using pAN456 (Nelson et al., 2007).

In the obtained transgenic line carrying *proPIN1:PIN1-mGFP4:PIN1term* (Benková et al., 2003), the insert was genetically linked with the *EPFL2* gene, and we were not able to outcross this marker into the *epfl1 epfl2* mutant. Using DNA from the transgenic plants, we reamplified *proPIN1:PIN1-mGFP4:PIN1term* and inserted it into pPZP222 between the BamHI and KpnI sites creating plasmid pAMO103.

To generate the proERL1:H2B-eGFP:35Sterm construct (pESH751), H2B-eGFP was amplified and inserted into pESH245 using the SalI and BamHI restriction sites. To generate the proERL2:H2B-eGFP:35Sterm construct (pESH752), *H2B-eGFP* was amplified and inserted into pESH246 using the SalI and PstI restriction sites. pESH245 and pESH246 vectors have been described previously (Shpak et al., 2004). They contain the *GUS* gene under the control of either *ERL1* or *ERL2* promoter in the pPZP222 backbone.

The inserted sequences in pAMO100, pAMO101, pAMO102b, pAMO103, pAMO105, pMIG101, pESH751, and pESH752 plasmids were confirmed by Sanger sequencing. The full sequence of pAMO119 and pAMO124 plasmids was determined by Oxford Nanopore Technology at Plasmidsaurus. Plasmids were transformed into an *Agrobacterium tumefaciens* strain GV3101/pMP90 by electroporation and introduced into Arabidopsis plants by the simplified floral dip method (Clough and Bent, 1998). pPZK411, pPZK412, pAMO100, pAMO101, pAMO105 and pMIG101 were introduced into the *epfl1 epfl2* mutant. pAMO102b was introduced into *epfl1 epfl2* and the wild type. pPZK401, pPZK402, and pAMO124 were introduced into the wild type. The *proCUC3:er-CFP* transgenic plants were previously described (Gonçalves et al., 2015). This marker was outcrossed into the *epfl1 epfl2* mutant. pAMO119 was introduced into the wild type and *epfl1 epfl2* plants both containing *proCUC3:er-CFP* (Gonçalves et al., 2015). The plasmid with dexamethasone-inducible EPFL2 under EPFL3 promoter (pAMO113) has been described earlier (Uzair et al., 2024). This plasmid was transformed into the *epfl1 epfl2* mutant. pAMO103 was at first introduced into the *epfl1 epfl2* mutant. Later, this marker was outcrossed from the mutant into the wild type using one stable transgenic line. pESH751 and pESH752 were introduced into a Landsberg erecta ecotype that has a point mutation K_750_→I_750_ in the *ERECTA* gene (*er-1* allele) (Torii et al., 1996). In T1 and T2, all transgenic plants described above, except pPZK401 and pPZK402, were selected using 100 mg/L and 60 mg/L gentamycin, respectively. Transgenic plants expressing pPZK401 and pPZK402 were selected using 25 mg/L kanamycin. Multiple homozygous T3 lines were generated for each construct.

The *pGreenIIM proDR5v2:ntdTomato/proDR5:n3GFP* and pGreenIIM proRPS5A:mDII-ntdTomato/proRPS5A:DII-n3Venus plasmids were described previously (Liao et al., 2015) and obtained from AddGene (plasmids #61628 and #61629, respectively). They were transformed into the *epfl1 epfl2* mutant. Transgenic plants were selected using MS plates with 0.1 mg/L methotrexate, and multiple homozygous T3 lines were generated. *proDR5v2:ntdTomato/proDR5:n3GFP* or *proRPS5A:mDII-ntdTomato/proRPS5A:DII-n3Venus* markers were outcrossed from one selected *epfl1 epfl2* line into the wild type.

Wild-type seeds expressing pGREENII proARF5:SV40-3x eGFP (CS67076) were obtained from the ABRC stock center and outcrossed into the *epfl1 epfl2* mutant. The proCUC2:CUC2-GFP construct (also called CUC2g-GFP) (Temman et al., 2023) was outcrossed from the wild type into the *epfl1 epfl2* mutant. The *AtML1pro:ER-FLAG* construct (Uchida et al., 2012) was outcrossed from the *er-105* mutant into *epfl1 epfl2*. *proWUS:H2B-eGFP* marker (Zhang et al., 2021) was outcrossed from the wild-type into *epfl1 epfl2*.

### Microscopy

Inflorescence clusters containing flowers up to stage 12 were harvested from the primary inflorescences after a plant made 10-15 flowers. They were placed in 90% ethanol 10% acetic acid solution to rock overnight and then gradually rehydrated by replacing the solution with 90% ethanol, 70% ethanol, 50% ethanol, 20% ethanol, and water with 20 minutes of rocking in between each step. Afterward, the samples were placed in 0.5 M NaOH solution and incubated at 37^0^ C for 15 to 20 minutes to make them transparent, then rinsed with water several times to remove residual NaOH before Differential Interference Contrast (DIC) microscopy. To measure ovule density, stage 10 flowers were identified based on developmental markers such as the presence of all ovule primordia in a finger-like shape prior to integument initiation and the early differentiation of the style (Smyth et al., 1990; Schneitz et al., 1995; Yu et al., 2020). Aniline blue staining followed the above tissue-clearing protocol described above. After the final rinsing step, the sample was placed in 0.1% (w/v) aniline blue solution in 50mM potassium phosphate pH 7.5 buffer for 30 minutes to one hour in a protocol adapted from (Lu, 2011). The samples were then rinsed with potassium phosphate buffer and imaged using a Nikon Eclipse 80i epifluorescence microscope with a 12-megapixel cooled color camera and a UV-2A filter (Nikon). Ovules in the process of meiosis II were identified by the presence of two strong callose bands stained by aniline blue, as previously described (Cao et al., 2018). Ovules that were fully straightened out on the slide and in focus were imaged. Ovule height was measured using ImageJ by drawing a line from the middle of the funiculus’s base to the nucellus’s tip on calibrated images.

For confocal microscopy, inflorescence clusters containing flowers up to stage 12 were harvested and fixed in 4% paraformaldehyde under a vacuum for up to 2 hours. Samples were then rinsed with 1x Phosphate-buffered saline (PBS) pH7.4 twice before placing them in ClearSee solution consisting of 25% (w/v) urea, 15% (w/v) sodium deoxycholate, and 10% (w/v) xylitol as previously described (Kurihara et al., 2015). Microcentrifuge tubes with samples were wrapped with foil to prevent photobleaching and rocked for 1 to 2 days with a daily change of ClearSee solution. Leaving samples in the solution longer than 2 days will lead to the loss of the signal. The day before imaging, samples were placed in a staining solution containing 0.1% (v/v) SCRI Renaissance 2200 (SR2200) dye, 1% (v/v) DMSO, 0.05% (v/v) Triton-X100, 5% (v/v) glycerol in ClearSee solution and left to rock overnight (Musielak et al., 2015). Before imaging, the samples were rinsed with fresh ClearSee. Water instead of ClearSee was used as a mounting medium to prevent contamination of the microscope water immersion lens. Slight dissection separated and exposed stage 8 through stage 10 flowers. Confocal microscopy was performed using a Leica SP8 Laser Confocal microscope. Samples were imaged using a 40x water immersion lens at 1024 x 512-pixel size with a zoom factor between 1.0x to 1.5x. Bidirectional scanning was used with a scan speed between 200 and 600 Hz, and line averaging was used with a line average of 3. Z-stacks were generated with a Z-step size of no greater than 0.5 microns.

The SR2200 dye was excited with a 405 nm solid-state UV laser and imaged using a 420-450 nm emission window. CFP was excited with a 456 nm argon laser and imaged using either a 460-500 nm emission window or, if GFP was also expressed in the sample, with a 460-480 nm emission window. GFP was excited with a 488 nm argon laser and imaged using a 500-520 nm emission window. Red fluorescent proteins such as mCherry or tdTomato were excited with a white light laser at 556 nm and imaged using a 605-625 nm emission window. Sequential line scanning was used to image multiple channels at once. Hybrid detectors were used when available, while PMT detectors were occasionally used when necessary. Laser power, gain, and emission filters were modified depending on the brightness of the signal and the number of fluorophores present in the sample. Confocal z-stacks were processed into 2D images using FIJI software. For each channel, background subtraction and contrast enhancement were used if necessary to remove background and enhance signal-to-background ratios. For longitudinal cross sections of ovule initiation, the slices of the z-stack that represent the center of the ovule primordia, were identified. Next, a Z-projection with frame averaging was created using 3 to 5 slices around the center of ovule primordia to create a smoother image while not exceeding the width of an individual cell. Finally, images were cropped to the desired size in microns, and image size in pixels was readjusted so that all images in an individual figure had the same dimensions in both pixels and microns.

### Chemical Treatments of Inflorescences

To study the effect of gibberellins and auxin on ovule initiation, inflorescence clusters were briefly dipped into a water solution that contained 0.01% Tween-20, 0.05% DMSO, and one of the following: 25 μM gibberellic acid (GA), 4 μM paclobutrazol (PAC), 10 μM N-1-naphthylphthalamic acid (NPA) or 50 μM 2,4-Dichlorophenoxyacetic acid (2,4-D). The mock solution contained 0.01% Tween-20 and 0.05% DMSO. Dipping was done five days in a row, once a day at the same time each day. Inflorescence clusters were collected on day six, and ovule initiation was studied using DIC microscopy. To study the effect of cytokinins on ovule initiation, inflorescence clusters were dipped into a water solution that contained 0.01% Silwet L-77, 0.1% DMSO, and 1 mM 6-benzylaminopurine (BAP). The mock solution contained 0.01% Silwet L-77 and 0.1% DMSO. Inflorescence clusters were dipped three days in a row and collected on day four.

## Acknowledgments

We thank Mitsuhiro Aida for sharing with us seeds of *cuc2-3* and *cuc3-105* mutants and *proCUC2:CUC2-GFP* transgenic plants. We thank Nicolas Arnaud for *proCUC3:er-CFP* transgenic plants. We thank Andreas Nebenfuehr for the pAN456 plasmid and Zackary Nimchuk for the pENT H2BGFP plasmid. We thank Keiko Torii for seeds of *er-105* with *AtML1pro:ER-FLAG*. We thank Andreas Nebenführ and Albrecht von Arnim for commenting on the manuscript.

Accession Numbers: BZR1 (At1g75080), CUC2 (At5g53950), CUC3 (At1g76420), DRN (At1g12980), EPFL1 (At5g10310), EPFL2 (At4g37810), EPFL4 (At4g14723), EPFL6 (At2g30370), ERECTA (At2g26330), ERL1 (At5g62230), ERL2 (At5g07180), PIN1 (At1g73590), WUS (At2g17950)

## Supplemental Data

The following supplemental materials are available.

**Figure S1.** EPFL1 and EPFL2 promote ovule initiation by positively regulating ovule number.

**Figure S2.** EPFL1, EPFL2, EPFL4, and EPFL6 synergistically regulate placenta elongation.

**Figure S3.** Auxin is able to accumulate into maxima during ovule initiation in the *epfl1 epfl2* mutant.

**Figure S4.** The efflux transporter of auxin PIN1 is expressed similarly in the wild type and *epfl1,2* mutant.

**Figure S5.** Inhibition of auxin transport severely reduces ovule density, while exogenous auxin does not affect either placenta length or ovule initiation.

**Figure S6.** Exogenous cytokinins promote placenta elongation, while exogenous gibberellins inhibit ovule density.

**Figure S7.** EPFL1, EPFL2, CUC2, and CUC3 redundantly regulate the length of the placenta.

**Figure S8.** Expression of *EPFL2* and CUC2 is not altered in *cuc2,3* and *epfl1,2*mutants, respectively.

**Figure S9.** During ovule outgrowth, *DRN* and *CUC3 are* expressed longer next to each other in the *epfl1 epfl2* mutant compared to the wild type.

**Figure S10.** The *bzr1-1D* mutation inhibits ovule initiation in the *epfl1,2* background.

**Figure S11.** The expression of *DRN* is expanded in the *epfl1 epfl2* mutant compared to the wild type.

